# Improving the sensitivity of cluster-based statistics for fMRI data

**DOI:** 10.1101/2020.01.06.896167

**Authors:** Linda Geerligs, Eric Maris

## Abstract

Because of the high dimensionality of neuroimaging data, identifying a statistical test that is both valid and maximally sensitive is an important challenge. Here, we present a combination of two approaches for fMRI data analysis that together result in substantial improvements of the sensitivity of cluster-based statistics. The first approach is to create novel cluster definitions that are sensitive to physiologically plausible effect patterns. The second is to adopt a new approach to combine test statistics with different sensitivity profiles, which we call the min(p) method. These innovations are made possible by using the randomization inference framework. In this paper, we report on a set of simulations that demonstrate (1) that the proposed methods control the false-alarm rate, (2) that the sensitivity profiles of cluster-based test statistics vary depending on the cluster defining thresholds and cluster definitions, and (3) that the min(p) method for combining these test statistics results in a drastic increase of sensitivity (up to five-fold), compared to existing fMRI analysis methods. This increase in sensitivity is not at the expense of the spatial specificity of the inference.

## Introduction

Functional magnetic resonance imaging (fMRI) is a tool that is widely used for clinical and basic neuroscience. The statistical analysis of fMRI data is mostly performed in a parametric framework, using cluster-based statistics. Recent work by Eklund, Nichols, and Knutsson (2016) has shown that false alarm (FA) rate control in within-subjects functional magnetic resonance imaging (fMRI) analyses using cluster based statistics was less accurate than expected, especially for low cluster defining thresholds (CDTs, e.g. p<0.01). This has reinforced the use of strict CDTs to maintain accurate FA rate control. The downside of this practice is a substantial reduction in sensitivity (statistical power), especially for detecting widespread but weak effects.

The failure to detect true activations due to lack of statistical power leads to low reproducibility of the results. In the current scientific climate, this may be an even more pressing problem than poor FA-rate control (Button et al., 2013; Szucs & Ioannidis, 2017). One way to improve statistical power is by increasing the study sample sizes, ideally motivated by a formal power analysis. However, despite an increase in the number of studies with large datasets, the median sample sizes in fMRI studies were still below 30 in 2015 (Poldrack et al., 2017). Here, we present an alternative way to improve the statistical power of fMRI studies: we will demonstrate that the sensitivity of statistical tests can be substantially increased (up to five-fold) by combining two approaches: (1) creating test statistics that are sensitive for physiologically plausible effect patterns, and (2) adopting a new approach for combining test statistics with different sensitivity profiles.

To achieve this goal, we operate within the randomization inference framework. This framework has a number of important advantages over parametric frameworks for fMRI analysis because it allows (1) to prove FA rate control under a relevant null hypothesis (statistical independence between the biological data and the explanatory variable; see further), (2) the use of an arbitrary test statistic, which allows us to select a test statistic solely on the basis of its sensitivity to the effects of interest, and (3) to combine test statistics with different sensitivity profiles (e.g., different CDTs). All of these advantages will be illustrated by the simulations on which we report in this paper.

The present paper builds on a theoretical paper on randomization testing in fMRI data (Maris, 2019). This theoretical paper provides the formal proof of FA rate control and presents a discussion on how to design a study such that it can be statistically analyzed under the randomization framework.

In the remainder of this paper we will first provide a recipe for how the randomization test can be used for a within-participant study and discuss ways to optimally design a test statistic. Next, we will use a set of simulations to demonstrate accurate FA rate control and illustrate that the use of different test statistics can substantially increase the sensitivity of fMRI data analysis.

## Methods

### Data

Resting state fMRI data from 103 healthy controls from Oulu dataset in the 1000 Functional Connectomes Project (Biswal et al., 2010) were used for all analyses (http://fcon_1000.projects.nitrc.org/fcpClassic/FcpTable.html). We used the Oulu dataset because previous work showed poor FA-rate control for the permutation tests with this particular dataset (Eklund et al., 2016). The dataset includes 37 male and 66 female participants with a narrow age range (20-23 years, mean=21.52, SD=0.57). Collection of the data was approved by the ethics committee of the Northern Ostrobothnian Hospital District. Data were collected using a 1.5 Tesla MR scanner, with a repetition time (TR) of 1.8 seconds. The data consist of 245 time points per subject, 64 × 64 × 28 voxels of size 4 × 4 × 4.4 mm.

### Simulation design

The aim of the simulations was to investigate the FA-rate control and the sensitivity of different test statistics within the randomization framework in comparison to the current standards in the field. We focused on group-level analyses comparing two different task conditions. The resting state data were used as the background signal on top of which we added a simulated stimulus-evoked signal. When the expected magnitude of the stimulus-evoked signal was equal in the two task conditions, we could investigate the FA-rate control of different statistical tests. To investigate the sensitivity profiles of the different statistical tests, we manipulated (1) the between-condition difference in the expected magnitudes of the evoked signals (denoted as “effect size” in the following), and (2) the size of the grey-matter volume that exhibited this difference (denoted as “spatial extent” in the following). The effect sizes were quantified as the (population-level) Cohen’s d of the between-condition differences in the signal magnitudes. The values of Cohen’s d in our simulation design were 0 (for investigating the FA-rate control), 0.6, 0.8, 1 and 1.2. The task-related BOLD signals were added to the resting state data (see *Simulating Data*) in a cluster of voxels which was defined by a sphere with a radius of 10, 15 or 20 mm centered at MNI coordinate [3 −60 30] (see Figure 1). Thus, our simulation design was 5 (effect size) by 3 (spatial extent).

**Figure 1:**
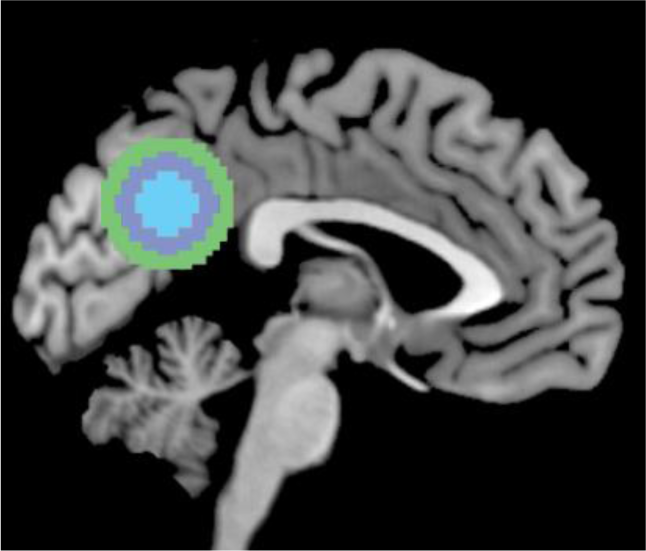
illustration of the sizes of the (unsmoothed) simulated clusters with a radius of 10, 15 or 20 mm.

### Simulating data

For each of the 15 cells of our simulation design, we performed 1000 group analyses, each of which started from a random sample of 40 participants from the Oulu dataset. These 1000 random samples of participants were the same as in Eklund et al. (2016) and were kept constant across all statistical tests.

We simulated an event-related paradigm with two stimulus sequences, of which one was assigned to condition A and the other to condition B. Each stimulus sequence had 62 simulated stimulus onsets with random durations (1-4 seconds) and inter-stimulus intervals (3-6 secs; same as E2 in Eklund et al., 2016). The order of stimulus durations and inter-stimulus intervals was reversed for sequence 2 as compared to sequence 1. The same stimulus sequences were used for all participants. In line with the argument in *A recipe for performing a randomization test for a within-participants study*, we simulated data for two complementary condition orders. For one condition order, one sequence was associated with condition A and the other to condition B. For the complementary condition order this was reversed. In each random sample of 40 participants, half of the participants were randomly assigned to one condition order and the other half to the complementary order.

The amplitude of the simulated evoked blood-oxygen-level-dependent (BOLD) response varied across participants but was the same for different events within the same individual. For each participant in the Oulu dataset, the amplitude of the evoked response in each of the two task conditions was drawn from a normal distribution with a mean of 4.5 and SD of 2.25. Using these values, when contrasting each task condition with the baseline, a significant cluster was found in approximately 90% of 1000 stimulations (using GRF-based inference with a cluster defining threshold of p<0.001). Differences in the evoked responses between task conditions were introduced by adding a constant value to the evoked response amplitudes of all participants for condition A. The size of this constant was chosen such that the effect size of the between-condition difference for the evoked response amplitudes (quantified as Cohen’s d) was either zero (for testing the FA-rate control), 0.6, 0.8, 1 or 1.2.

Two task-related BOLD signals were created for each participant by convolving the stimulus functions (specifying onset and duration) with the canonical hemodynamic response function (HRF) and multiplying the resulting time course with the amplitude of the evoked response in each task condition. After convolution with the canonical HRF, the regressors for conditions A and B were uncorrelated (r<0.01).

The task-related BOLD signals were added to the resting state data in a cluster of voxels which was defined by a sphere with a radius of 10, 15 or 20 mm centered at MNI coordinate [3 −60 30] (see Figure 1). To ensure that the assumptions of GRF theory were met, the image containing the cluster definition (specified as zeros and ones) was smoothed with a Gaussian kernel (6 mm FWHM, same as the smoothness of the resting state data, see *fMRI data analyses*). When combining the task-related BOLD signal and the resting state data, the task signal was multiplied by the weights in the cluster definition image. True positive voxels were defined as voxels that were part of the cluster definition before smoothing. True negative voxels were defined as voxels that contained less than 1% of the task-related signal (i.e., after smoothing). Only these true positive and true negative voxels were considered in metrics of sensitivity or specificity of the statistical tests.

### fMRI data analyses

The fMRI data were pre-processed using standard SPM 8 processing pipelines (http://www.fil.ion.ucl.ac.uk/spm/software/spm8/), including realignment, co-registration, normalization and 6 mm FWHM smoothing (for more details, see Eklund et al., 2016). The fMRI data were not corrected for geometric distortions, as no field maps are available.

A general linear model (GLM) was applied to the preprocessed fMRI data, using two regressors for each of the two task conditions (A and B): the HRF-convolved stimulus function (specifying onsets and durations) and its first derivative. The stimulus onset and duration times in the GLM were matched to the stimulus onset and duration times that were used to simulate the data (which depend on the condition order). The estimated head motion parameters were used as additional regressors in the design matrix, to reduce effects of head motion. To account for low-frequency drifts in the data, a discrete cosine transform with cutoff of 128 seconds was used. Temporal correlations were corrected for with a global AR(1) model in SPM. The first-level contrast between task conditions A and B (A-B) was used as the input for the group-level analyses.

In the group-analyses, we looked for voxels or clusters in which the A-B contrast was significantly different from zero. As a baseline for the evaluation of the performance of the randomization test, we investigated the FA rate and the sensitivity of the following alternative statistical methods: thresholding using false discovery rate control (FDR, Genovese, Lazar, & Nichols, 2002), cluster-level Gaussian random field (GRF) inference (Friston, Worsley, Frackowiak, Mazziotta, & Evans, 1994), and control using the randomization distribution of threshold-free cluster enhancement (Smith & Nichols, 2009). For the analyses relying on the randomization distribution, we used the maximum value the test statistic of interest (summed within-cluster t-statistics or voxel-specific TFCE values) to correct for multiple comparisons.

## Results

### A recipe for performing a randomization test for a within-participants study

The randomization inference framework we propose here, tests the null hypothesis of statistical independence between the biological (i.e. fMRI data) and the explanatory variable (i.e., the experimental conditions). Statistical independence involves that, for the biological data of a randomly sampled participant, it does not matter in which experimental condition it is observed.

The randomization test relies on randomization of the explanatory variable across participants. For a within-participant study, the explanatory variable is the order in which the conditions are presented (denoted as “condition order” in the following). The randomization inference framework requires that there are multiple condition orders that reflect the effect of interest. To clarify the steps that are involved in performing the randomization test, we give an example for one specific study (see Figure 2). This example study involves eight trials and two experimental conditions (A and B), and 20 participants are completing both experimental conditions. Importantly, the first step occurs prior to the data collection.

**Figure 2:**
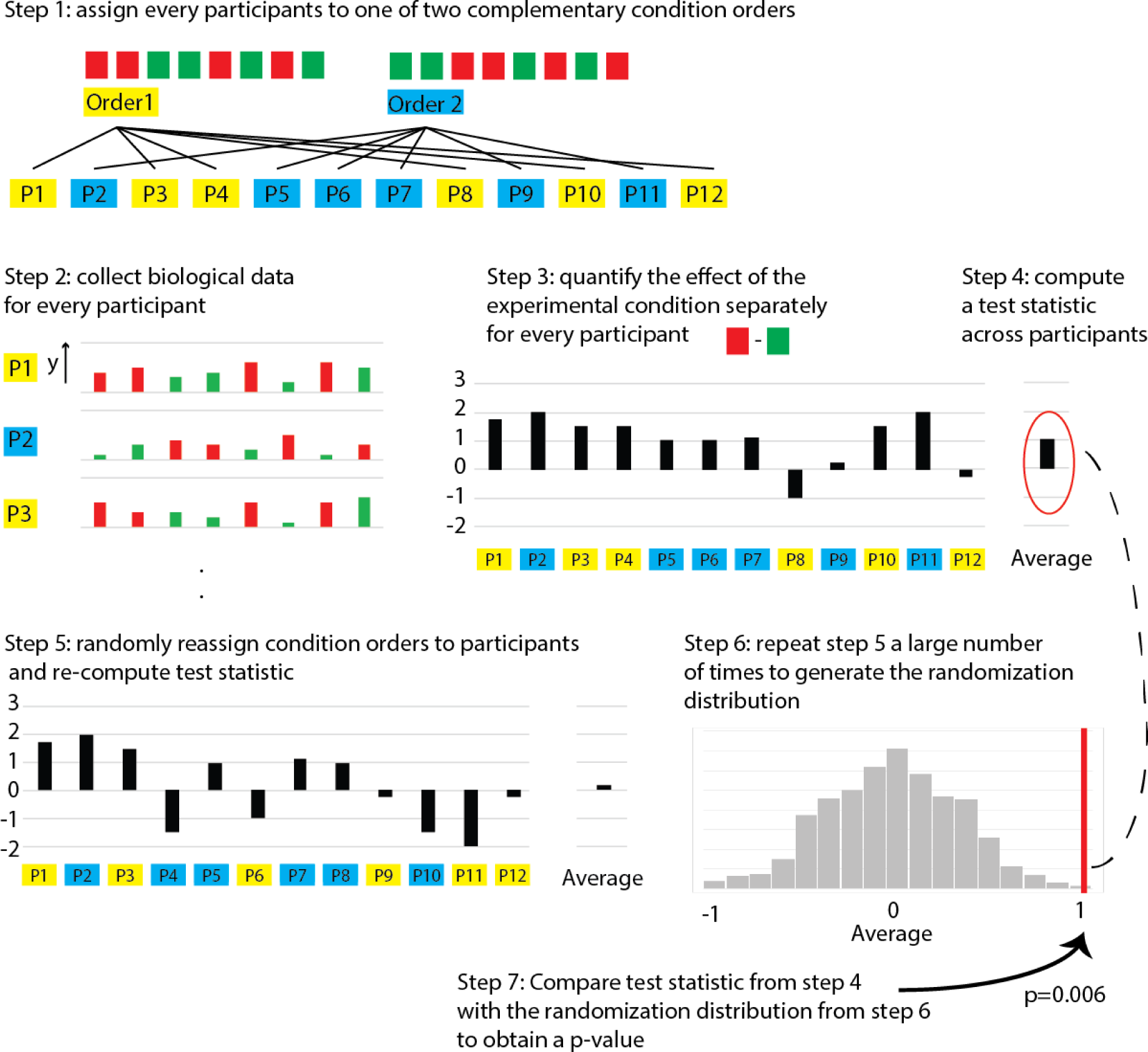
Schematic explanation of the steps in the randomization test. This example represents the procedure for a test with a single variable of interest (i.e. one voxel), but the same steps apply in the case of cluster-based statistics.

1. The participants are randomly assigned to one of two condition orders. To optimize sensitivity, it is important to select condition orders that are as different as possible, for example, [AABBABAB, BBAABABA].
2. The fMRI data is collected for every participant.
3. The effect of the experimental conditions (A versus B) is quantified separately within every participant. When analysing fMRI data, this is commonly done by running a multivariate regression, and representing conditions A and B as separate regressors. Contrast images, which reflect the difference between the beta values of the regressors A and B are typically the basis for the quantification of the effect of interest.
4. A test statistic is computed by combining the effects identified in step 3 across participants. Typically, this is done by computing a t-statistic across the contrast images. However, the randomization framework allows for any other statistic that reflects the difference between conditions A and B. In the case of cluster-based statistics, clusters are typically identified by applying a cluster-defining threshold (CDT) and then counting the number of voxels in this cluster or summing its thresholded voxel-level statistics. Usually, multiple clusters are identified, and the test statistic is then taken as the maximum (for thresholding from below) or the minimum (for thresholding from above) of the cluster-level statistics. The randomization framework allows for many variations on this typical way of calculating cluster-based statistics (see section ‘How to construct a test statistic?’).

5+6. The randomization p-value is calculated for the observed test statistic. This is done using a reference distribution that is obtained by randomly reassigning the participants to one of the two condition orders, while keeping the observed data (i.e. the single subject contrast images) fixed. Participants are randomly reassigned to one of the two condition orders and the maximum/minimum cluster-based statistic is recalculated. Repeating these steps (random re-assignment and recalculation) a large number of times results in the randomization distribution, which is the reference distribution of a randomization test.

7. Each of the observed cluster-level statistics can be compared to the reference distribution to obtain a randomization p-value. If one of these p-values (the smallest one, which corresponds to the maximum/minimum observed cluster-based statistic) is less than the nominal alpha level, the null hypothesis of statistical independence between the biological data and the explanatory variable is rejected.

The example above illustrates the simplest scenario for the randomization test. In some cases, it is not possible to randomly assign participants to condition orders before the data collection. For example, when the explanatory variable is not under experimental control (e.g., behavioural outcome, non-BOLD physiological variables like EEG and pupil diameter) or when analyzing an existing dataset in which more than two condition orders were used. In those cases, it is very often possible to use the randomization framework by relying on prior knowledge or weak assumptions about the probability distribution of the explanatory variable. For example, consider an existing dataset in which participants were randomly assigned to one of *all possible* condition orders. For this scenario, a valid and sensitive randomization test is also obtained if the randomization distribution is constructed by randomly re-assigning every participant to either (1) the condition order to which they were actually assigned, or (2) the complement of that particular participant-specific condition order. Thus, every participant has its own pair of complementary condition orders, of which one member is always the observed condition order. If the test statistic is some function of the t-statistics across the contrast images, then the random reassignment either maintains or flips the sign of the test statistic. This random reassignment procedure is only possible if one knows the probabilities of the original random assignment. The FA rate control of this random reassignment procedure is proved in Maris (2019), which also describes how to deal with explanatory variables that are not under experimental control.

### How to construct a test statistic?

One essential advantage of the randomization framework proposed here, is that it allows the researcher to construct a test statistic that is maximally sensitive to the effects of interest. There are many different ways a test statistic for cluster-based inference can be constructed. The first consideration is which voxel-connectivity structure should be used. The connectivity structure determines which voxels should be treated as each other’s neighbors, thereby defining the basis for merging voxels in a cluster. Connectivity between voxels can be defined as voxels that share a corner with the current voxel (26 neighbors for each voxel – C26, the FSL default), voxels that share an edge (18 neighbors – C18, the default in SPM) or voxels that share a surface (6 per voxel – C6, the default in AFNI, see Figure 3A). A stricter connectivity definition will result in smaller sized clusters, and this affects the sensitivity for detecting a cluster with some shape of interest. We propose defining neighbors as voxels that are connected via a surface (C6). This will reduce the chance of identifying clusters with voxels that are only connect through a series of corners or edges, as we believe that such a narrow thread is biologically implausible. In our simulation study (see further), we compared the sensitivity of cluster-based test statistics that involve different connectivity definitions.

**Figure 3:**
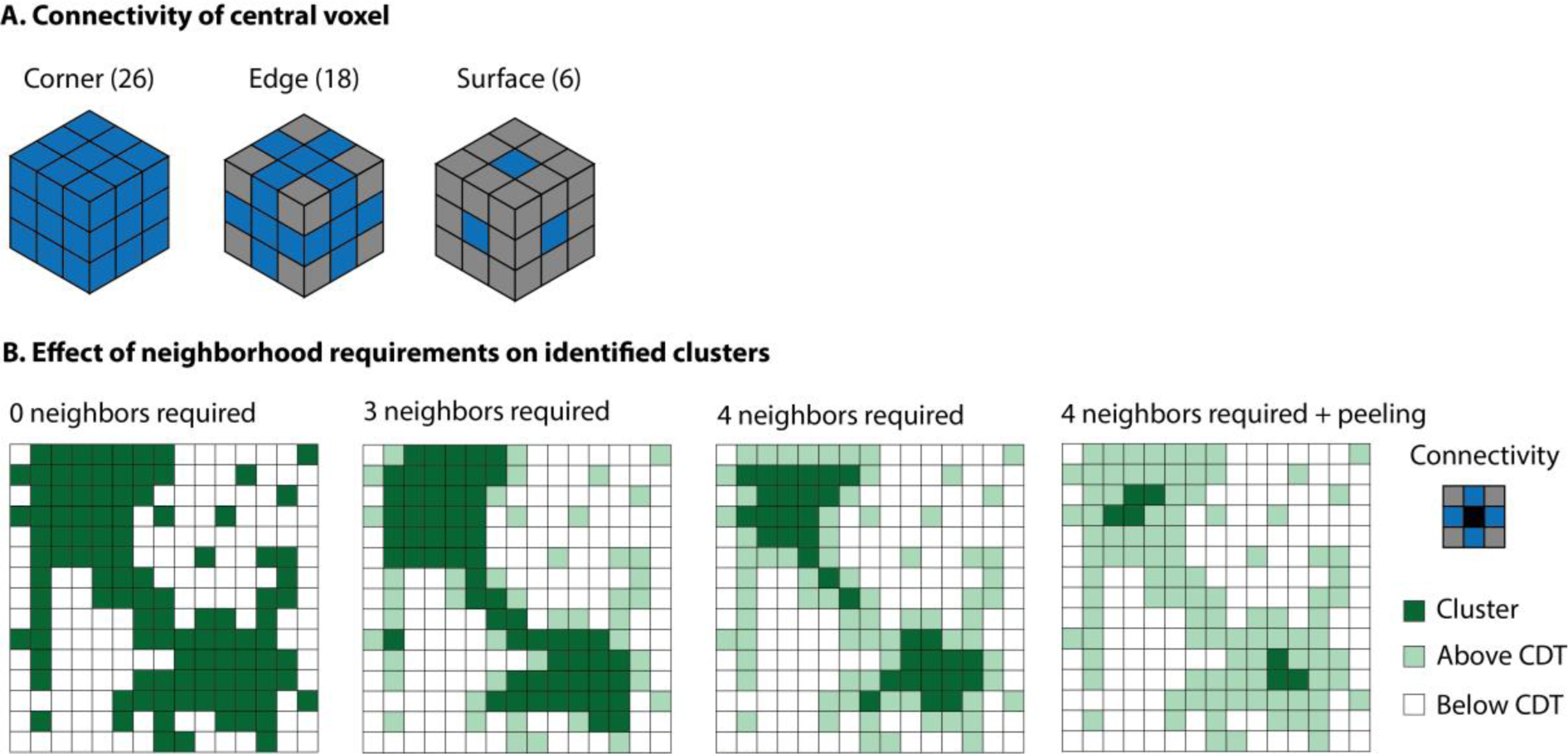
(A) Illustration of the three different voxel connectivity structures. Blue voxels are neighbors of the central voxel. (B) 2D illustration of the effect of neighborhood requirements on the identified clusters.

Another way to vary the cluster definition is by varying the CDT. Strict CDTs are best suited to detect large effects that are present in a small number of voxels. Lenient CDTs are best suited to detect small effects that are present across a large area of the brain. There are also different ways to quantify to the magnitude of a cluster. In the parametric framework, a cluster’s magnitude is typically quantified by its size: the number of voxels within the cluster. However, other quantifications such as the sum over the t-values of within-cluster voxels have been shown to be a more sensitive measure in EEG data (Maris & Oostenveld, 2007). Here we use the sum over t-values to determine cluster magnitude.

To devise a potentially sensitive test statistic for fMRI data, we also looked at the cluster patterns in a large number of spatial maps of thresholded t-statistics that were calculated on data without an effect. We observed that especially at low cluster defining thresholds, there were often two or more separate clusters that were connected via a narrow thread of voxels in between. This resulted in larger cluster sizes in the randomization distribution and less sensitive statistical tests. To avoid such sprawling clusters, it is possible to adapt the cluster definition. One option is to impose a minimum number of above-threshold neighbors that each voxel should have before it is included in the cluster. Here, were we will investigate cluster definitions with no restrictions on the minimum number of neighbors (N0), at least 3 neighbors (N3), 5 neighbors (N5) or 6 neighbors (N6). By removing voxels with a low number of above-threshold neighbors, it is possible to counteract the effects of the data smoothness that is often introduced during preprocessing: voxels at the edge of the cluster will be removed while voxels at the center remain. Single voxels or very small clusters of above threshold voxels are biologically implausible and can be the result of spatial smearing around isolated voxels with high t-values. The effects of these restrictions on the number of neighbors is illustrated in Figure 3B. To further reduce such sprawling clusters, instead of a single removal of voxels with a low number of above threshold neighbors, we can perform this removal several times (denoted as “iterative peeling”, with shorthand notation P#, in which the # denotes the number of iterative removals minus one). This way, we can avoid both clusters of isolated voxels, as well as clusters with a small volume that may haphazardly merge to form a larger sprawling cluster. Table 1 provides an overview of the cluster definitions we examined in this paper and the shorthand we will use to refer to those definitions in the remainder of the paper. Each of these cluster definitions can be paired with different cluster defining thresholds (CDTs).

**Table 1:** Overview of cluster definitions

| Shorthand | Connectivity structure (C) | Neighborhood requirements | Peeling |
| --- | --- | --- | --- |
| C18N0P0 | 18, sharing edges | None | No |
| C6N0P0 | 6, sharing surfaces | None | No |
| C6N3P0 | 6, sharing surfaces | Min. 3 active neighbors | No |
| C6N5P0 | 6, sharing surfaces | Min. 5 active neighbors | No |
| C6N6P0 | 6, sharing surfaces | Min. 6 active neighbors | No |
| C6N6P1 | 6, sharing surfaces | Min. 6 active neighbors | Apply neighborhood definition in two iterations |

### How to combine different test statistics?

A crucial advantage of the randomization framework is that it allows to combine cluster statistics with different sensitivity profiles. A researcher may not know whether to expect small or large clusters. In that case, it is possible to analyze the data using different cluster definitions, for example by varying the CDT, the neighbor definition, and/or the number of required neighboring voxels. Within the randomization framework, the results can be combined over these different cluster definitions. For each of the different cluster definitions (CDT, neighbor definition, etc.), the randomization step results in a distribution of optimum (i.e., maximum or minimum) cluster magnitudes (size or sum). These randomized optimum cluster magnitudes can each be transformed into p-values by comparing them to their corresponding randomization distribution (see Figure 4). By definition, and separately for each of the cluster definitions, the probability distribution of these p-values is uniform (see Figure 4, Step 2). Similarly, each observed cluster magnitude can also be transformed into a p-value by comparing it to its corresponding randomization distribution. After transforming the cluster-definition-specific magnitudes into p-values, these transformed magnitudes can be meaningfully combined in a single randomization distribution. This is realized by taking the minimum p-value over all cluster definitions. This min(p) randomization distribution is constructed in a loop over draws from the randomization distribution: for every draw, evaluate which of the cluster definitions (statistics) has the smallest p-value, and use the resulting value as a realization of the min(p) randomization distribution. This min(p) randomization distribution is the final distribution that is used for decision making: if the observed min(p)-value is less than the *α* × 100-th percentile of the min(p) randomization distribution, then we reject the null hypothesis of statistical independence between biological data and the explanatory variable. By using the min(p) randomization distribution for decision making (instead of the cluster-definition-specific randomization distributions), we correct for multiple testing (one test per cluster definition).

**Figure 4:**
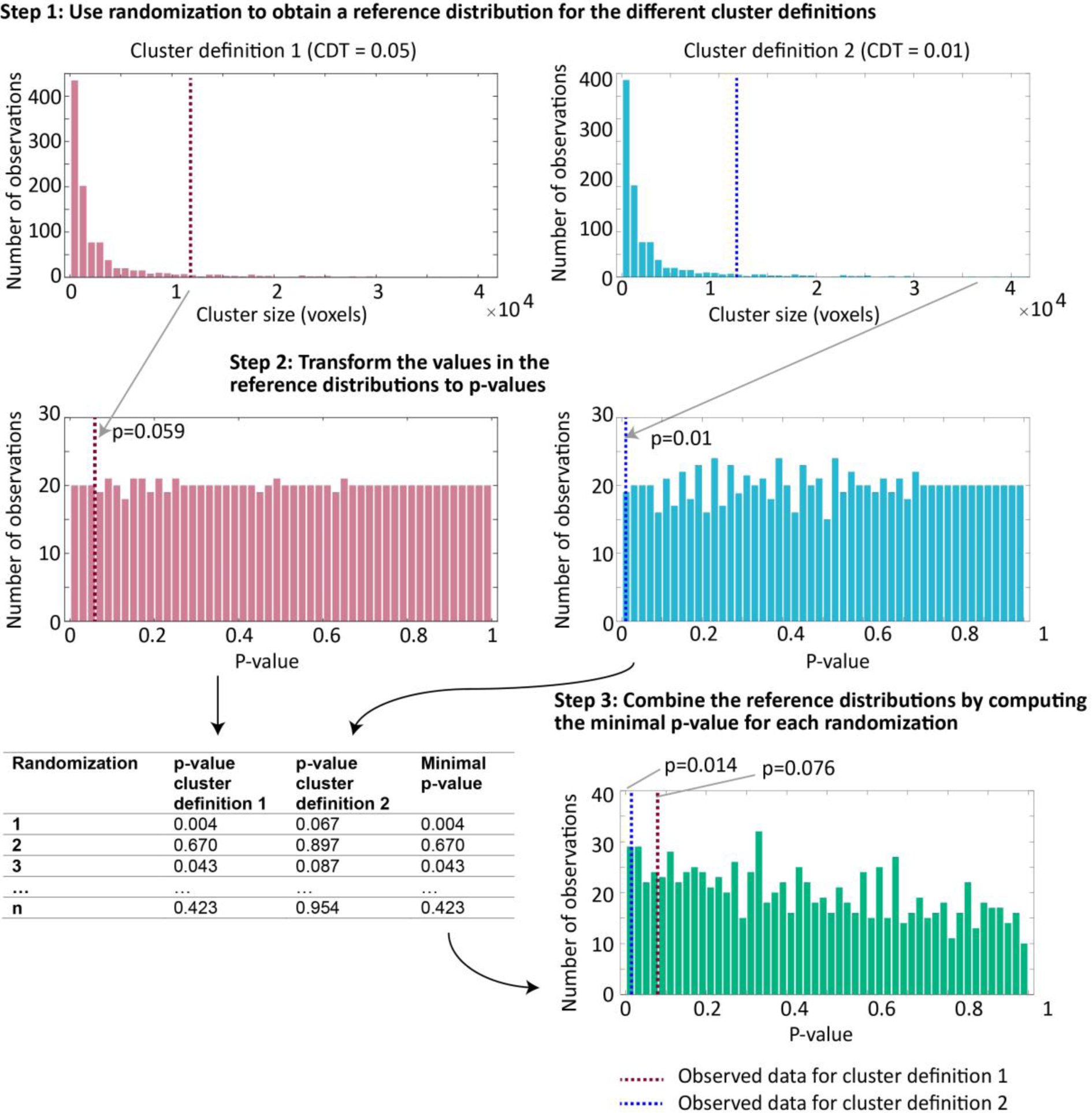
Illustration of how to combine cluster statistics with different sensitivity profiles. This illustration is for combining cluster definitions with different CDTs, but the method is the same for combining other test statistics or more than two test statistics.

### False alarm rate control

The randomization tests on which we report in this paper all control the FA rate under a highly relevant null hypothesis: statistical independence between the biological data and the explanatory variable. The explanatory variable that we are considering here are the condition orders to which the participants are randomly assigned (in the example above, AABBABAB or BBAABABA). In neuroimaging, researchers typically interpret their effects in terms of amplitude of the stimulus-evoked hemodynamic response (HR) in relation to occurrence of particular stimuli. At first sight, this differs from our null hypothesis, which is formulated in terms of the whole biological data, as recorded. However, it can be demonstrated that our whole-data-level null hypothesis is implied by a null hypothesis at the level of the stimulus-evoked HR amplitude: statistical independence between the stimulus-evoked HR amplitude and the explanatory variable (Maris, 2019). Therefore, if the latter HR-level null hypothesis is false, then so is the null hypothesis at the level of the whole biological data.

The formal proof of the FA rate control of our randomization test is presented in a companion paper (Maris, 2019). For the present paper, the crucial aspect of this proof is that it holds for all possible test statistics. This creates the opportunity to construct test statistics that optimize the sensitivity for a wide range of profiles (weak and widespread, strong and focal, etc.). The essence of the proof is that, under the null hypothesis, (1) conditionally on the biological data, the probability distribution of the test statistic only depends on the randomization distribution (the distribution that assigns the participants to the condition orders), and (2) conditional FA rate control implies unconditional FA rate control. In its simplest form, this proof applies to randomized experiments only. However, by introducing the concept of “conditioning on an informative support” the proof can be extended to studies in which the explanatory variable passively observed (e.g., disease status, accuracy, response time, EEG power) instead of under experimental control.

As a part of our simulation study, we empirically checked the correctness of the proof in Maris (2019). Different from the noise-only simulation studies by Eklund et al. (2016), we simulated fMRI data in which every participant’s data exhibited nonzero stimulus-evoked HR amplitudes to both experimental conditions within a restricted cluster of voxels (see methods). Because our interest was in FA rate control, the expected values of these stimulus-evoked HR amplitudes (calculated over the population of participants) were equal in the two conditions (see Methods). We calculated the FA-rates of the cluster definitions in Table 1, each combined with four CDTs (0.05, 0.01, 0.005 and 0.001), and compared these with the FA-rates of three alternative popular methods: false discovery rate control (FDR; Genovese et al., 2002), cluster-level Gaussian random field (GRF) inference (Friston et al., 1994), and control using the randomization distribution of threshold-free cluster enhancement (TFCE; Smith & Nichols, 2009). FDR and GRF control have a rationale in the parametric framework, and TFCE achieves FA rate control by making use of the randomization distribution of the maximum TFCE-value (instead of the maximum cluster statistic, as in our approach). TFCE can be performed for different connectivity structures, and here we used the surface (TFCE-C6) and the corner connectivity structure (TFCE-C26). TFCE depends on two tuning parameters, and for the first set of results we used the values that were also used in the original publication of the method (Smith & Nichols, 2009). In a later set of results (*Combining TFCE-parameters*), we also report on the performance of TFCE with other parameter values.

The results of our simulation study in Figure 5 supported the proof in Maris (2019): the randomization test controlled the FA-rate for each of the cluster definitions in Table 1 and each of the CDTs. FDR and the randomization-based TFCE also controlled the FA rate, although FDR was too conservative. Importantly, for the parametric inference based on Gaussian random-field theory (GRF) the FA-rate was only accurately controlled for a CDT equal to 0.001; the FA-rate rose up to 32% for parametric inference with a CDT of 0.05. Therefore, in the remainder of the paper, we only consider GRF-based inference using a CDT equal to 0.001.

**Figure 5:**
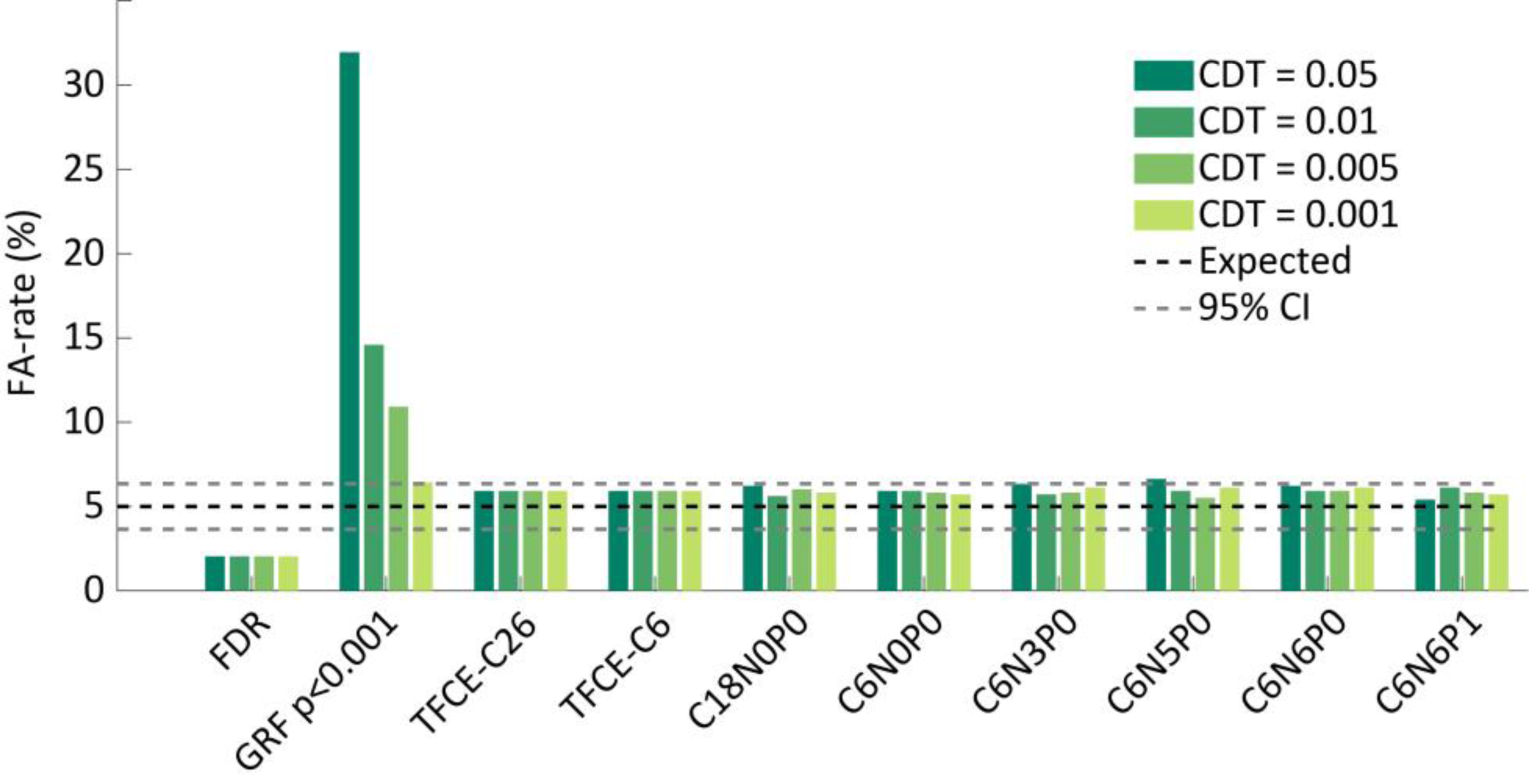
The observed FA-rate for the Gaussian random field theory (GRF) and for each of the six different cluster definitions that were used in the simulations. The dotted lines show the binomial 95% confidence interval around the 5% nominal FA-rate for 1000 simulations.

### Sensitivity

The main objective of our simulation study was to investigate the sensitivity of the different test statistics to detect plausible simulated effects in the data. To this end, we simulated data using stimulus-evoked HR amplitudes of which the expected values differed between the two conditions (see Methods). In the simulation design, we varied across four effect sizes (Cohen’s d’s of 0.6, 0.8, 1 and 1.2; see Methods) and three simulated true cluster sizes (spheres with radia of 10, 15 and 20 mm; see Methods). As a baseline for our comparisons, we calculated the sensitivity of the three alternative popular methods: FDR, GRF and randomization-based TFCE.

There are several ways of quantifying sensitivity. Here, we quantify sensitivity as the proportion of simulations in which the significant clusters (those with a p-value less than 0.05) cover at least half of the simulated cluster (called *identification* rate). This reflects our interest in also identifying the location of the effect, instead of only detecting the presence of an effect somewhere in the brain. The latter aspect can better be quantified as the proportion of simulations in which there are significant clusters (called *hit* rate). Later, and for comparison, we will also show results for this latter aspect of sensitivity.

Figure 6 shows that, when the cluster is large or the effect size is small, TFCE shows the highest identification rate. In the other cases, GRF with CDT=0.001 is the most sensitive test. FDR is the least sensitive test statistic. TFCE with the surface connectivity structure (TFCE-C6) is slightly more sensitive than TFCE with the corner connectivity structure (TFCE-C26). Therefore, we will use TFCE-C6 and GRF with CDT=0.001 as the reference statistics in the comparison with our cluster statistics.

**Figure 6:**
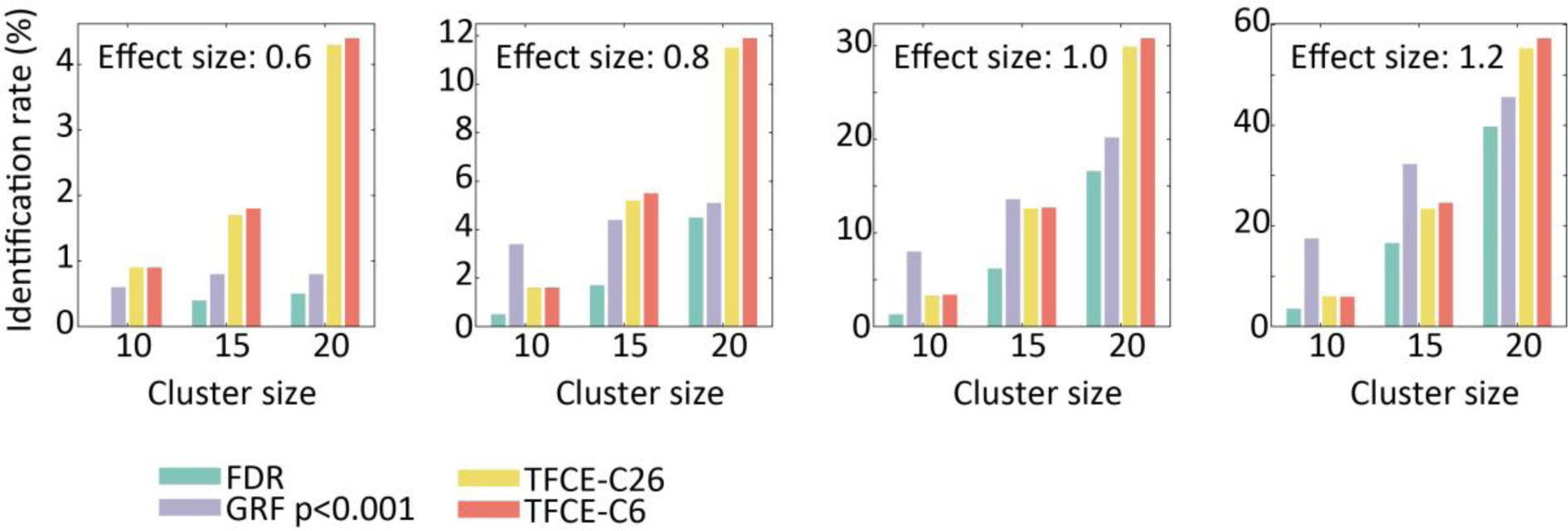
The observed identification rates for four test statistics that are commonly used in fMRI research. The identification rate is the % of simulations in which a significant cluster overlaps with at least half of the simulated cluster.

Next, we compared the sensitivity of the different cluster definitions in Table 1 across the four CDTs. Figure 7 shows that the choice for a particular CDT and cluster-definition has a large impact on the sensitivity of the statistical test. We observed that the sensitivity of the test statistics involves a trade-off between CDT and cluster definition: cluster definitions with more neighborhood restrictions tend to be more sensitive when they are combined with more lenient CDTs, while cluster definitions with fewer restrictions tend to be more sensitive when combined with stricter CDTs. Also, for small effect sizes, the lenient CDTs are always more sensitive than the strict CDTs. On the other hand, for intermediate and large effect sizes, the strict CDTs tend to be more sensitive in the case of a small cluster size while the more lenient CDTs tend to be more sensitive for a large cluster size. The three cluster definitions with the fewest constraints on neighborhood structure (C18N0P0, C6N0P0 and C6N3P0) showed very similar sensitivity patterns, and therefore we will only consider C6N3P0 in the remainder of the paper.

**Figure 7:**
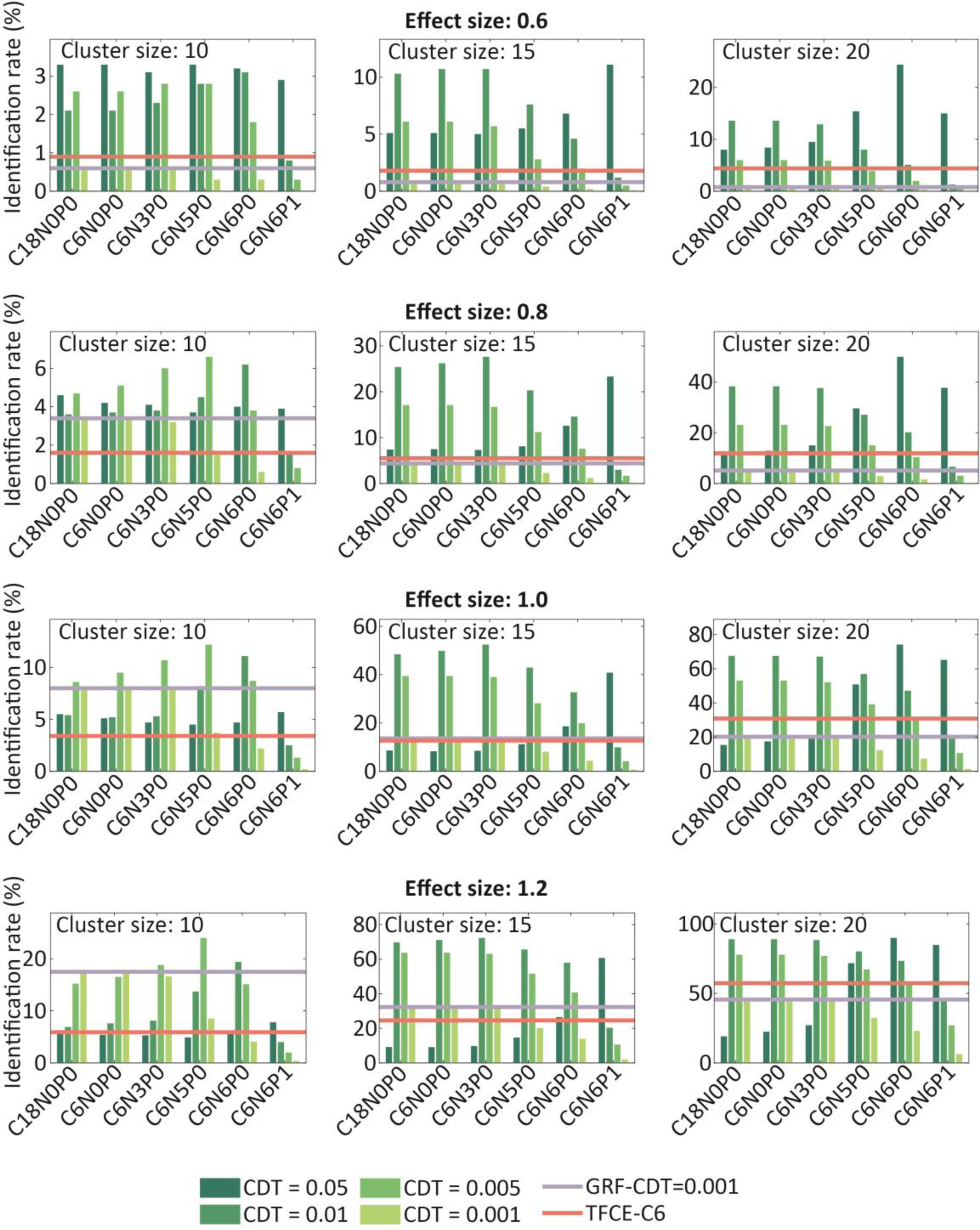
The observed hit rates for each of the basic different cluster definitions that we used in the randomization testing framework. As a reference, we also plotted the identification rates for GRF with a CDT of p<0.001 and for TFCE-C6.

Crucially, we find that for all effect sizes and cluster sizes, we can identify at least one test statistic that outperforms both GRF at p<0.001 and TFCE-C6. However, the sensitivity to different effect sizes and cluster sizes differs widely across test statistics (i.e., cluster-definition-CDT combinations) and it is not possible to choose a single test statistic that performs best in all cases.

### Combining cluster definitions

Figure 7 illustrates that different test statistic are optimal for different combinations of effect sizes and cluster sizes. This was motivation for combining different test statistics by means of the min(p) method (see the section *How to combine different test statistics?*). In particular, for each of the four remaining cluster definitions (C6N3P0, C6N5P0, C6N6P0 and C6N6P1) we computed a combined test statistic that combines across different CDTs (0.05, 0.01, 0.005 and 0.001). We also used the min(p) method to compute a combined cluster statistic that combines across all of these four cluster definitions and CDTs. The combined test statistics for the four cluster definitions (across all CDTs) are denoted as minp-C6N3P0, minp-C6N5P0, minp-C6N6P0 and minp-C6N6P1, and the combined test statistic across all cluster definitions and CDTs is denoted as minp-all.

Figure 8A illustrates that combining the C6N3P0 test statistic over CDTs using the min(p) method results in a test statistic that has a sensitivity that is similar to the best performing CDT, for all cluster sizes and effect sizes. For large and small cluster sizes the minp-C6N3P0 statistic performs a little better than the best performing CDT, while for the intermediate cluster size it performs a little bit worse. Our finding that the minp-C6N3P0 statistic performs as well or nearly as well as the optimal CDT for the C6N3P0 statistic shows that the correction for multiple testing (i.e., the multiple CDT-specific C6N3P0 statistics) using the min(p) method has only minimal effects on the sensitivity.

**Figure 8:**
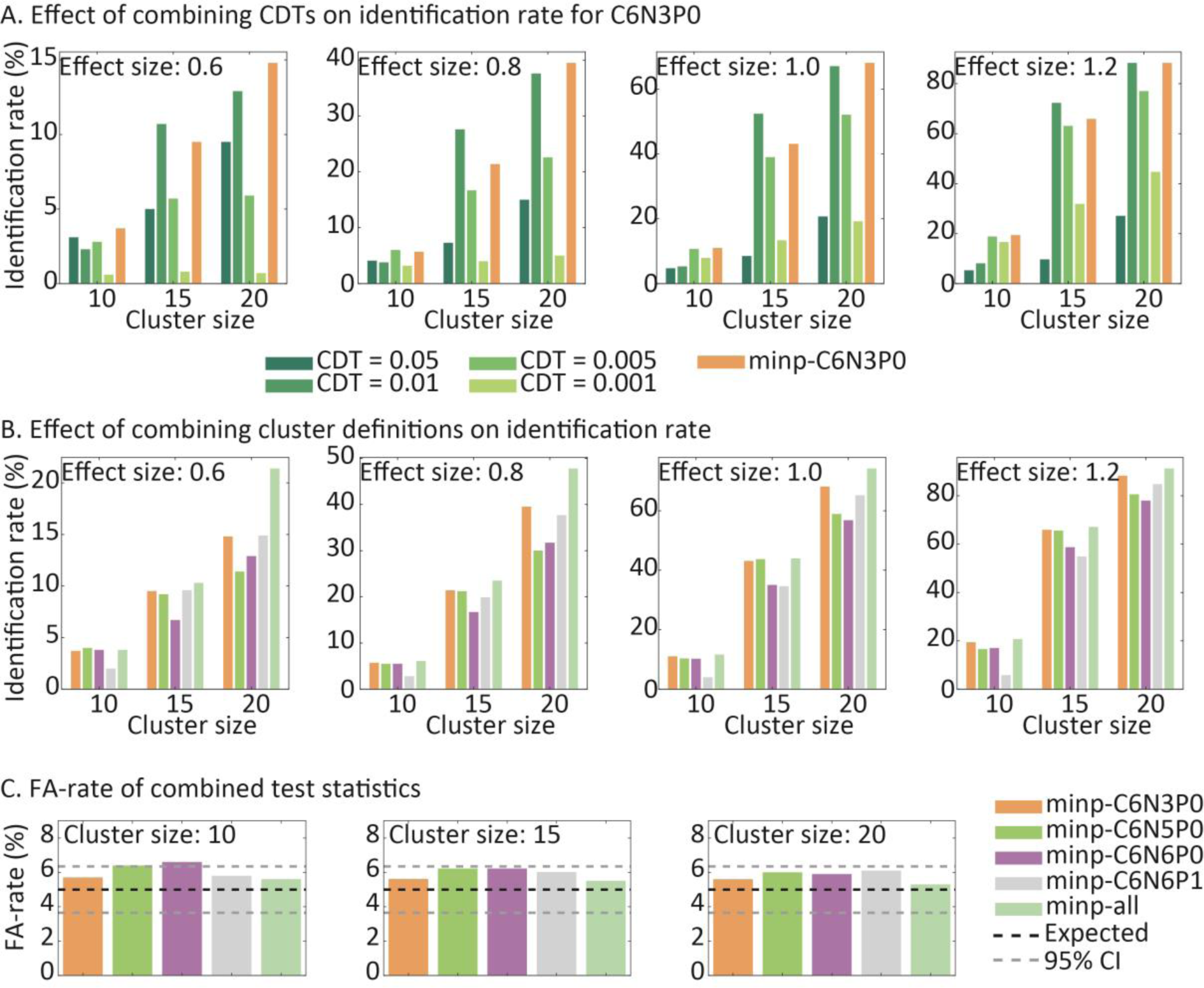
(A) The effect of combining CDT’s using the min(p) method on the identification rate is illustrated for the C6N3P0 test statistic. (B) The effects of combining cluster definitions using the min(p) method on the identification rate. Each of the cluster definitions was also combined across CDTs using the min(p) method. (C) FA-rates of the combined cluster definitions.

The results for the other three cluster definitions (C6N5P0, C6N6P0 and C6N6P1) are highly similar. However, there are substantial differences *between* the different cluster definitions, as is also clear from Figure 7. This fact motivates the use of the minp-all statistic, for which the results are shown in Figure 8B. This figure shows that combining different cluster definitions using the min(p) method results in further improvements in sensitivity. In fact, for all cluster sizes and effect sizes, the minp-all statistic is equally sensitive or more sensitive than the best performing single cluster definition statistic. This shows that combining different cluster definitions using the min(p) method results in a better sensitivity for the different types of effect.

As a further check on the proof in Maris (2019), we also calculated the empirical FA-rates for the different min(p) test statistics. Figure 8C shows that all these test statistics control the FA-rate at their nominal values.

### Combining TFCE-parameters

In Figure 8, we reported on the results that were obtained by combining different CDTs and different cluster definitions into one single test statistic by means of the min(p) method. The same method can also be used to combine across the different tuning parameter values for TFCE. The calculation of the TFCE image depends on a width and a height parameter, and different values for these parameters may result in a different sensitivity profile of the associated statistical test. Therefore, we also investigated the sensitivity of a min(p)-TFCE statistic that combined across all 25 tuning parameter combinations that were considered in the original paper by (Smith & Nichols, 2009). We applied this method to TFCE with surface (TFCE-C6) connectivity structure. Figure 9A shows that the minp-TFCE-C6 statistic shows better sensitivity than TFCE-C6 for small effect sizes. However, for intermediate and large effects sizes we did not observe an advantage of combining across TFCE parameter settings.

**Figure 9.**
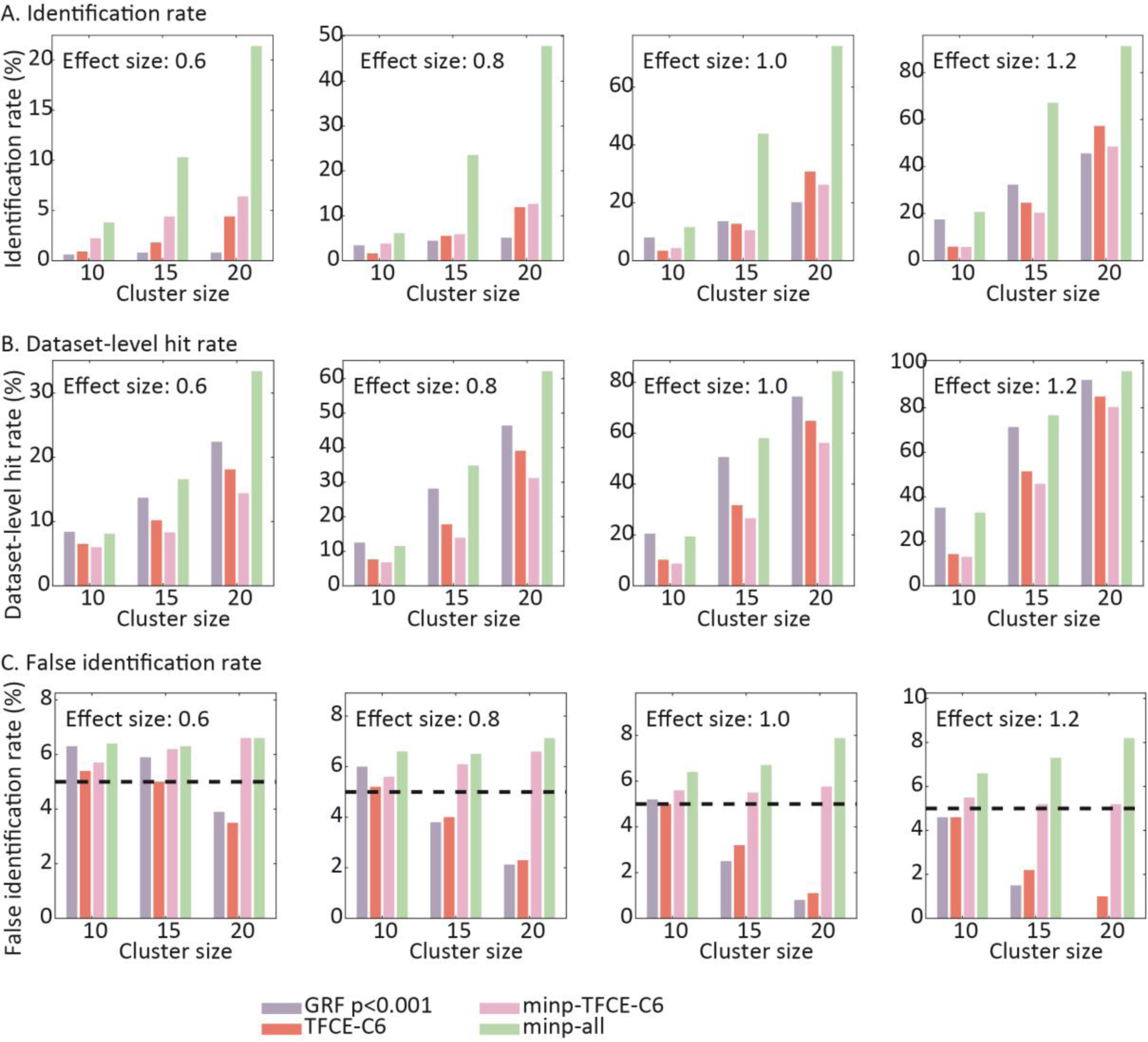
Different measures of sensitivity and spatial specificity of the GRF, TFCE, TFCE-MP and minp-all test statistics. (A) The identification rate, which is the % of simulations in which a significant cluster overlaps with at least half of the simulated cluster. (B) The dataset hit rate, which shows the percentage of simulations in which an above threshold cluster was detected, regardless of whether this overlapped with the simulated cluster. (C) The spatial specificity, which is measured as the percentage of simulations in which more false-positive than true-positive voxels were detected.

### Comparing the best test statistics

We now compare the four test statistics that are the most promising on the basis of our previous analyses: GRF with CDT=0.001, TFCE-C6, minp-TFCE, and minp-all. Figure 9A shows the identification rates for these four test statistics. We found that, for all simulated cluster sizes and effect sizes, minp-all outperformed all three other test statistics (the existing methods). For small effect sizes (0.6 and 0.8) and intermediate or large clusters, the identification rate of minp-all is up to 5 times larger than the one for the best performing existing method. For larger effect sizes (1.0 and 1.2), the identification rate increases for all test statistics, but minp-all continues to outperform the existing methods. For example, for large clusters with an effect size of 1, the identification rate increases from 31% to 74%.

Identification rate, the measure of sensitivity on which we reported so far, is the probability that at least 50% of the voxels in the simulated cluster is detected as significant. This measure depends on two factors: (1) the probability that one or more clusters are significant, and (2) the probability that these significant cluster cover more than 50% of the voxels in the simulated cluster. Because of this second factor, the identification rate is also a measure of effect coverage. An alternative measure of sensitivity reflects only the first factor: the probability that at least one significant cluster is identified, irrespective of whether or how much this activation overlaps with the simulated cluster. We refer to this measure of sensitivity as the dataset-level hit rate, and we show the results in Figure 9B. In terms of the dataset-level hit rate, for medium and large clusters, the minp-all statistic outperformed all other test statistics. For small clusters, however, GRF at CDT=0.001 slightly outperformed the minp-all statistic. Together, the results in Figures 9A and B suggest that GRF at CDT=0.001 is good at detecting whether there is an effect, but performs poorly in identifying the spatial extent of the effect. The minp-all statistic performs well for both measures of sensitivity.

The flipside of a better effect coverage is a potentially reduced spatial specificity. To investigate whether this is indeed the case, we calculated the percentage of simulations in which the number of false positive voxels was larger than the number of true positive voxels, which we will call the *false identification rate*. The result of this analysis is shown in Figure 9C. For small cluster sizes, the false identification rate for all four test statistics is approximately the same. For intermediate and large clusters, the false identification rate is higher for the minp-all statistic than for the existing methods. However, in absolute terms, the minp-all statistic showed adequate spatial sensitivity for all cluster and effect sizes, as the false identification rate was always between 6 and 8%.

## Discussion

We have described a randomization test that can be used for a within-participant neuroimaging study. This randomization test controls the FA rate under the null hypothesis of statistical independence between the biological data and the explanatory variable (Maris, 2019). Because the FA rate control of a randomization test does not depend on the test statistic, we discuss ways to design a test statistic such that sensitivity is optimized. Specifically, we introduce the min(p) method for combining test statistics with different sensitivity profiles. We performed a set of simulations that demonstrate accurate FA rate control and illustrate the different sensitivity profiles of cluster-based test statistics with different CDTs and cluster definitions. Using the min(p) method for combining these test statistics resulted in a drastic increase of sensitivity, improving on the existing methods for statistical analysis of fMRI data. This increase in sensitivity was not at the expense of the spatial specificity of the inference.

This paper belongs to the long tradition of nonparametric statistical methods for the analysis of neuroimaging data (Bullmore et al., 1996; Hayasaka & Nichols, 2003; Maris & Oostenveld, 2007; Nichols & Holmes, 2002; Winkler, Ridgway, Webster, Smith, & Nichols, 2014). There are three essential differences between the existing literature and the present paper. First, the present paper builds on a formal framework for testing the novel null hypothesis of statistical independence between the biological data and the explanatory variable (Maris, 2019), whereas previous work has mainly focused on exchangeability and distributional symmetry (Maris & Oostenveld, 2007; Nichols & Holmes, 2002; Winkler et al., 2014). Because the null hypothesis is formulated at the level of the raw data (instead of functions of the raw data, such as, regression coefficients), this formal framework allows for a straightforward application to event-related designs. In contrast, null hypotheses about functions of the raw data require distributional assumptions that may be violated (Eklund et al., 2016; Winkler et al., 2014).

Second, the present paper introduces and applies the min(p) method for combining test statistics with different sensitivity profiles. The min(p) method is novel and is not exclusively linked to the null hypothesis of statistical independence and its associated formal framework. Specifically, the min(p) method can also be used in the framework that is based on exchangeability and distributional symmetry (Maris & Oostenveld, 2007; Nichols & Holmes, 2002; Winkler et al., 2014). In addition, it cannot only be applied to studies with a within-participant manipulation of the explanatory variable (the focus of the present paper), but also to studies with a between-participant manipulation, and to observational studies in which the explanatory variable is not under experimental control (and can vary both within- and between-participants). The idea of constructing test statistics with specific sensitivity profiles is not novel. In fact, it lies at the heart of the TFCE methodology (Smith & Nichols, 2009). However, it is often unknown which sensitivity profile is optimal for a given dataset, and the min(p) method effectively deals with this ignorance by combining the different sensitivity profiles.

Third, our main quantification of sensitivity involved a measure that indexes coverage probability (identification rate), instead of the usual dataset-level hit rate. This quantification is in line with the main scientific interest in current neuroimaging research: identifying the location of the neural tissue that is affected by some experimental contrast. Especially in terms of coverage probability, our best performing cluster statistic (minp-all) outperformed all the existing statistical methods. Importantly, our simulations show that spatial specificity did not appreciably suffer from combining different test statistics.

As with every simulation study, its results do not have the status of a mathematical proof; it cannot be excluded that different results may be obtained with other ingredients for the simulation study (e.g., effect topography, noise correlations, test statistics). In other words, its conclusions depend on the parameters that were manipulated and the ones that were kept constant. An important parameter that was kept constant in our simulation study is the shape of the simulated clusters (spherical).

Similar to the effects of varying cluster size and effect size, we expect that different cluster definitions will be differentially sensitive to different cluster shapes. By combining across cluster definitions, the min(p) method would allow researchers to be sensitive to different types of cluster shapes. However, an in-depth investigation of the effect of different cluster shapes on the sensitivity of different cluster definitions is beyond the scope of the present paper.

Another important constraint in our simulation study is that there was no heterogeneity (individual differences) in the effect topography across the population of participants. An increase in this spatial heterogeneity would result in a decrease in the sensitivity of all statistical tests on which we reported. However, it is as yet unknown to which extent the statistical tests differ in exactly how much this sensitivity decreases. The randomization framework has the important advantage that it allows for test statistics that explicitly take into account individual differences with respect to effect topography. Such an approach would not produce a single “significant” effect topography. Instead, it would be a statistical test of the spatially aspecific null hypothesis of statistical independence of which the sensitivity is robust to individual differences in effect topography.

Analytic flexibility is one of the main threats of reproducible neuroimaging research (Poldrack et al., 2017). This practice involves that, after collecting the data, the researcher analyses his data in several different ways, with each analysis pipeline typically culminating in a statistical test. Combining the min(p) method with preregistration provides a sensitivity-preserving solution for the FA rate inflation that results from this analytic flexibility. This FA rate inflation follows from two errors: (1) designing analysis pipelines after inspecting the patterns in the data, and (2) failing to correct for multiple testing. The simplest prevention against the first error is preregistration, and the simplest remedy for the second error is Bonferroni correction. However, Bonferroni correction does not take into account the statistical dependence (correlation) between the different test statistics, and this goes at the expense of statistical sensitivity. The min(p) method is very likely to be a sensitivity-preserving alternative for Bonferroni correction because the randomization distribution of the minimum p-value *does* take this statistical dependence into account.

In conclusion, the present paper describes two statistical innovations for neuroimaging studies: (1) a randomization test for within-participant studies, and (2) a method for combining test statistics with different sensitivity profiles. These two innovations allow for novel statistical tests that control the FA rate and drastically outperform the existing statistical tests with respect to sensitivity. Future research has to show (1) whether the formal framework used in the present paper can also be used for improving sensitivity in other study types (e.g., studies involving explanatory variables that are not under experimental control), and (2) to what degree the results of our simulation study depend on its specific ingredients.

## Acknowledgement

LG was supported by a Veni grant [451-16-013] from the Netherlands Organization for Scientific Research. We thank Anders Eklund for sharing the pre-processed SPM data with us.

## Notes

#### Summary of Updates

Corrected a few minor errors in the figures.

http://fcon_1000.projects.nitrc.org/fcpClassic/FcpTable.html

